# Engineering Bacterial Biomanufacturing: Characterization and Manipulation of *Sphingomonas sp.* LM7 Extracellular Polymers

**DOI:** 10.1101/2024.05.16.594401

**Authors:** Ellen W. van Wijngaarden, Alexandra G. Goetsch, Ilana L. Brito, David M. Hershey, Meredith N. Silberstein

**Author notes:** Phone: 607/255-5063.

## Abstract

Biologically produced materials are an attractive alternative to traditional materials such as metals and plastics and offer improved functionalities such as better biodegradability and biocompatibility. Polysaccharides are an example of a biologically produced materials that can have a range of chemical and physical properties including high stiffness to weight ratios and thermal stability. Biomanufactured bacterial polysaccharides can come with many advantages such as being non-toxic and are mechanically robust relative to proteins and lipids, which are also secreted by bacteria to generate a biofilm. One major goal in biomanufacturing is to produce quality material quickly and cost-effectively. Biomanufacturing offers additional benefits compared to traditional manufacturing including low resource investment and equipment requirements, providing an alternative to sourcing fossil fuel byproducts, and relatively low temperatures needed for production. However, many biologically produced materials require complex and lengthy purification processes before use. This paper 1) identifies the material properties of a novel polysaccharide, dubbed promonan, isolated from the extracellular polymeric substances of *Sphingomonas sp.* LM7; 2) demonstrates that these properties can be manipulated to suit specific applications; and 3) presents two alternative methods of processing to shorten purification time by more than 50% while maintaining comparable material.

## 1 Introduction

Researchers are increasingly looking towards biomanufacturing to find alternative methods of material production that eliminate using fossil fuel byproducts and have low resource investment, mild growth temperatures, and comparatively less equipment required than traditional materials. A 2023 joint report from the U.S. Departments of Energy, Agriculture, Commerce, Health and Human Services and the National Science Foundation outlined goals to apply biomanufacturing to revolutionize agriculture, healthcare and supply chains through reducing emissions and plastic waste and providing novel health therapies.^1^ Goals 1.1 and 1.2 of the aforementioned report focus specifically on developing low carbon intensity materials and building a circular economy for materials. Biomanufacturing can help achieve these goals by harnessing existing cellular machinery to produce desired materials. Progress in the field has been facilitated by recent advances in synthetic biology and genetic modification techniques such as modern sequencing technologies^2^ and molecular genetic tools. ^3^ However, in order to successfully replace traditional materials, biomanufactured materials must demonstrate satisfactory mechanical and chemical properties, as well as be easy to produce in terms of time and required resources.

Materials manufactured by plants, fungi and bacteria come with a broad range of physical and chemical properties, making them ideal for a wide array of applications. ^4^ Bacteria are the most genetically tractable of the three options, with a wide variety of applicable synthetic biology tools. ^5^ Bacterial cells naturally produce an array of biopolymers including polysaccharides, polyphosphates, extracelluar DNA, and proteins that make up an extracellular matrix, known as a biofilm.^6^ Secreted polysaccharides aid cells with adhesion, aggregation, and structure.^7^ Well known producers of bacterial polysaccharides include *Xanthamonas campestris*,^8^ *Pseudomonas aeruginosa* ^9^ and *Sphingomonas sp.*^10^ Polysaccharides can be isolated and purified from a biofilm using enzymes and filtration methods.^8,10^

Bacterial polysaccharides have gained attention for their innovative uses. Common applications where bacterial polysaccharides are used include food processing,^11^ healthcare,^6,12^ and the oil and gas industry.^13^ Xanthan and dextran are two examples of bacterial polymers that are used in medicine, and as additives in cosmetics, food, and packaging. ^14^ Specific examples include xanthan gum used as a thickening agent in chocolate, a water binding agent in meat, and an additive to shampoo for suspending insolubles. ^8^ Another family of bacterial polysaccharides, called sphingans, enhance oil recovery from high temperature and high salinity reservoirs when used in a technique known as polymer flooding. ^15^ The unique rheological properties of bacterial polysaccharides enable them to be mixed homogeneously with other compounds to act as a thickening agent, achieving high viscosities at low concentrations. Furthermore, high biodegradability and non-toxicity make these materials ideal for use in consumable products. Finally, high resistance to degradation from salt and/or heat (often stable up to 300*^◦^*C) make bacterial polysaccharides versatile and ideal for broad use.^16,17^

Various manufacturing processes have been developed for the production of bacterial polysaccharides. Extracellular polysaccharides are secreted polymers that can be extracted efficiently by centrifuging to separate cells from polysaccharide. Precipitation is commonly performed with an alcohol such as methanol, ethanol, or isopropanol. ^18,19^ Other methods include metal coordination, quaternary ammonium salt precipitation, and column chromatography. However, these methods require difficult to source reagents, accurate pH control and are time-consuming.^20^ Following isolation of the material, impurities are typically removed using a combination of enzymes and liquid-liquid extractions using solvents such as phenol and chloroform.^20^ The method of isolation and purification greatly impacts the time and cost of production, while also affecting the material yield and purity. Additional work is needed to understand how these techniques alter material properties.

Many bacteria can produce polysaccharides but not all bacterial strains are suited for biomanufacturing. The ideal bacteria produce a medium amount of material under reasonable temperatures, between 20*^◦^*C and 40*^◦^*C, enabling growth in a standard lab incubator.^21^ It is important to note that sometimes too much material or too high of a viscosity can make processing challenging within a bioreactor setup.^22^ The type of material and quantity can be altered genetically to facilitate biomanufacturing if the organism is genetically tractable. In addition to genetically adjusting overall yield, the material properties can be fine-tuned genetically.^23^ Polysaccharides produced by the *Sphingomonas*genus, known as sphingans, can provide a broad range of properties including molecular weight (*M_W_*), surface charge,^15,16,24^ and thermal stability. These structural characteristics provide insight into the diverse material properties of sphingans that make these polysaccharides ideal for biomanufacturing.

The genus *Sphingomonas* was first established in 1990 and has since been put to use in many applications. These bacteria are known for producing high *M_W_* polysaccharides such as welan gum, gellan gum, and diutan gum. ^16^ Sphingans serve as good thickening agents, similar to other biopolysaccharides, and can also be used for chelating, emulsifying, and stabilizing.^16^ Unique rheological properties and stability at high temperatures and salinity have made Sphingans popular for use in oil recovery.^15^ One example, welan gum, exhibits a storage modulus of approximately 2 Pa when hydrated in water at a weight to volume (w/v) ratio of only 0.175%.^25^ Another example, gellan gum, is commonly used as a food thickener and has a storage modulus that is strongly frequency dependent, ranging from 0.01 to 10 Pa at a concentration of 1% w/v in water. ^26^ Sphingans have also been recently applied in bioremediation due to their hydrophobicity, minimal nutrient requirements for production, and ability to degrade various pollutants. Furthermore, many *Sphingomonas* strains are genetically tractable, holding potential for tailor-made sphingans. ^27^

This study investigates the properties of a newly isolated material, named promonan, from a novel strain, *Sphingomonas sp.* LM7, to identify possible uses and potential for biomanufacturing. Specifically, we address the following research questions: (a) How do the material properties of promonan compare with similarly structured bacterial polysaccharides? (b) Can these properties be manipulated to suit specific applications? (c) Can the purification process be shortened to improve manufacturability?

## 2 Materials and Methods

### 2.1 Bacterial Culturing

A 10 mL Peptone Yeast Extract media (PYE: 0.2% w/v peptone w/v yeast extract) with 3% sucrose starter culture was innoculated with LM7 cells from a frozen glycerol stock and grown at 30*^◦^*C overnight on a shaker. 10 mL of starter culture was added to 1 L of PYE and 3% w/v sucrose solution. The culture was grown until approximately the end of day 4 at 18*^◦^*C with vigorous shaking. All cultures were grown until an optical density at 600 nm of 0.5 was reached.

### 2.2 Enzymatic Purification Process

The full enzymatic purification process spans 8 days, following 2 days of culturing time, as outlined in Figure 1. Following growth, the cultures were then centrifuged at 16,000 x g for 90 minutes to remove cells. The supernatant was transferred to new containers in 150 ml aliquots. 250 ml ethanol was added to each tube and incubated overnight at 4*^◦^*C to perform cold ethanol precipitation. On day 5, the tubes were centrifuged at 16 k x g for 1 hour to result in the polysaccharide adhering to the bottom of the tube. The supernantant was decanted and the remaining pellet was left to dry for 30 minutes with the tube inverted. Each pellet was then resuspended in 7.5 ml deionized H_2_O. The solution was then transferred to a 50 ml falcon tube so that each falcon tube has a total volume of 15 ml.

**Figure 1:**
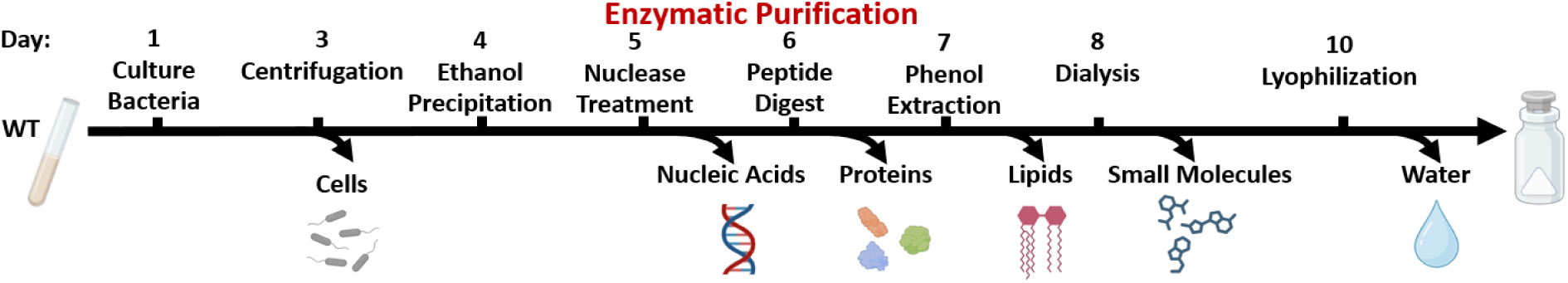
Timeline of the 10 day biomanufacturing process including enzymatic purification.

To perform nuclease digestion, 1 M Tris buffer and 1 M MgCl_2_ were added to the tubes to yield a final concentration of 10 mM and 2 mM respectively. Benzonuclease (Turbonuclease) (Accelagen Inc., San Diego, CA) was added at a concentration of 1 µl ml^-1^ to the tubes. The tubes were inverted and incubated at 37*^◦^*C overnight.

A peptide digest was performed by adding proteinase K at a concentration of 10 mg ml^-1^. Tris-HCl was added to reach a final concentration of 25 mM. The tubes were inverted and incubated at 37*^◦^*C overnight.

Phenol-chloroform extraction was performed by dividing volumes in half, 18 ml phenol and 2 ml chloroform were added to each 20 ml volume. The tubes were gently mixed and left to sit for 30 minutes to allow layers to separate. The upper aqueous layer was removed and tubes were combined in a new 50 ml falcon tube. The solution was then added to pre-wetted dialysis tubing with a molecular weight cutoff (MWCO) of 100 kDa.

Dialysis was conducted in a 4 L large beaker in deionized H_2_O. The water was changed daily for 3 days. Following dialysis, the solutions were pipetted in equal volumes not exceeding 4.5 ml into clean 11 ml glass tubes. The tubes were frozen at –20*^◦^*C for 2 hours before being transferred to the –80*^◦^*C freezer overnight. The caps were replaced with parafilm with a hole punched out or a piece of tissue for lyophilization. Samples were left in the lyophilizer (FreeZone 2.5 L Benchtop Freeze dryer, Labconco, Kansas City, MO) for 24 hours or until they were fully dry.

### 2.3 Dialysis-Based Alternative Biomanufacturing Process

The dialysis-based biomanufacturing process spanned 3 days, less than half the time required for the production process using enzymatic purification. After initial culture growth, the cells were removed via centrifugation as described in the enzymatic purification process. The solutions were then directly dialyzed for 24 hours, with 4 water changes approximately 6 hours apart (MWCO = 3.5, 50 or 100 kDa) and lyophilized for 24 hours.

### 2.4 Ultracentrifugation

Ultracentrifugation was used as an alternative physical purification method to investigate the effect of removing outer membrane vesicles, which usually contain proteins. Samples were centrifuged at 24,000 rotations per minute using an S-28 rotor at 4*^◦^*C in an Optima XL-100K Ultracentrifuge (Beckman Coulter, Brea, CA). The supernatant was decanted, frozen and lyophilized to isolate the material.

### 2.5 Protein Knockout Biomanufacturing Process

The protein knockout biomanufacturing process consisted of culturing the strain Δ*fliC*, removing cells as per the previously described enzymatic purification processes and then lyophilizing the material.

#### 2.5.1 Targeted Gene Deletion

Strains used in this study are listed in Supporting Table 1. The *fliC* gene (BXU08_RS13585) was deleted using a previously described method. ^27^ A construct that fused 491 base pairs (bp) upstream of the *fliC* open reading frame (ORF), the first twelve nucleotides of the *fliC* ORF, the last twelve nucleotides of the *fliC* ORF, and 508 bp downstream of the *fliC* ORF was designed. This DNA fragment was synthesized commercially (Azenta, Burlington, MA) and inserted into the SpeI/HindIII site of pNTPS138 to generate pDH1329. pDH1329 can generate an in-frame deletion, leaving a short protein coding sequence composed of the first and last four codons of the *fliC* open reading frame.

Wild-type LM7 or the Δ*prmJ* mutant were transformed with pDH1329 by electroporation, and primary integrants were identified by selection on PYE plates supplemented with 25 *µ*g mL^-1^ kanamycin. Colonies were restreaked on PYE plates containing 25 *µ*g mL^-1^ kanamycin and used to inoculate a liquid culture in PYE that was grown overnight under non-selective conditions. The outgrowth was then spread on PYE plates supplemented with 3% (w/v) sucrose to select for plasmid excision. Individual colonies were replica plated to confirm sensitivity to kanamycin and KanS colonies were screened using PCR to identify *fliC* deletion mutants.

#### 2.5.2 Preparation of Secreted Protein Extracts

Analyzing secreted proteins from wild-type LM7 cultures proved challenging because the high viscosity of extracts containing promonan interfered with efficient separation on SDS-PAGE gels. We instead used a mutant background (Δ*prmJ*) that cannot produce promonan to analyze secreted proteins. Overnight cultures of LM7 strains grown in PYE, were used to inoculate M2 medium supplemented with 2% (w/v) D-glucose. Cultures were grown at 18*^◦^*C for 48 hours with shaking at 200 rpm. Cells were removed from the culture broth by centrifugation at 3700 x g for 1 hour and 10 mL fractions of the resulting supernatant were transferred to fresh tubes. The spent medium extract was immediately added to 8 kDa cutoff dialysis tubing and dialyzed for 4 days against deionized water before lyophilizing until fully dry.

Dried spent medium extracts were resuspended in 250 *µ*L 2X SDS-LOAD buffer, heated at 70*^◦^*C for 15 minutes and centrifuged to remove insoluble material. Samples were loaded on polyacrylamide (10% w/v) gels, separated using the Tris-glycine buffer system and stained with Coomassie Brilliant Blue R. Individual bands were excised with clean razor blades and transferred to fresh tubes for protein identification.

#### 2.5.3 In-Gel Digestion of Extracted Proteins

Coomassie stained gel pieces were de-stained completely in MeOH/H_2_O/NH_4_HCO_3_ (50% / 50% / 25 mM) and dehydrated for 5 minutes in acetonitrile (ACN)/H_2_O/NH_4_HCO_3_ (50% / 50% / 25 mM) followed by 30 seconds in 100% ACN. Pieces were dried in a Speed-Vac for 1 minute, reduced in 25 mM Dithiotreitol in 25 mM NH_4_HCO_3_ for 15 minutes at 56*^◦^*C, alkylated with 55 mM CAA (Chloroacetamide in 25 mM NH_4_HCO_3_) in darkness at room temperature for 15 minutes, washed once in H_2_O, dehydrated for 2 minutes in ACN/H_2_O/NH_4_HCO_3_ (50% / 50% / 25 mM) then placed, once more, for 30 seconds in 100% ACN. The samples were dried again and rehydrated with 20 *µ*l of trypsin solution with 0.01% ProteaseMAX^TM^ surfactant comprised of 10 ng *µ*l^-1^ Trypsin (Promega Corp., Madison, WI) in 25 mM NH_3_HCO_3_ / 0.01% w/v of ProteaseMAX^TM^ from Promega Corp. The pieces were left to stand for 2 minutes at room temperature before adding 30 *µ*l of overlay solution (25 mM NH_4_HCO_3_/0.01% w/v of ProteaseMAX^TM^) to keep gel pieces immersed throughout the digestion. The digestion was conducted for 3 hours at 42*^◦^*C. Peptides generated from the digestion were transferred to a new tube and acidified with 2.5% TFA (trifluoroacetic Acid) to 0.3% final concentration. Gel pieces were additionally extracted with ACN:H_2_O:TFA (70% / 29.25% / 0.75%) for 10 minutes while vortexing. Solutions were combined and dried completely in a Speed-Vac (*∼*15 minutes). Extracted peptides were solubilized in 30 *µ*l of 0.5% TFA. Degraded ProteaseMAX^TM^ was removed via centrifugation (max. speed, 10 minutes) and the peptide solid phase extracted using 10 *µ*l volume C18 tips (*Pierce*^○R^ (Thermo Fisher Scientific, Waltham, MA) according to manufacturer protocol. Peptides were eluted off of the C18 SPE column with 5 *µ*l of ACN/H^2^O/TFA (70% / 30% / 0.1%), dried to completion, and resolubilized in 25 *µ*l total volume with 0.1% formic acid. A volume of 2 *µ*l was loaded on the instrument.

#### 2.5.4 Analysis of Tryptic Fragments by Nanoscale Liquid Chromatography tandem Mass Spectrometry (NanoLC-MS/MS)

Peptides were analyzed by nanoLC-MS/MS using the Agilent 1100 nanoflow system (Agilent Technologies, Santa Clara, CA) connected to a hybrid linear ion trap-orbitrap mass spectrometer (LTQ-Orbitrap Elite^TM^, Thermo Fisher Scientific) and equipped with an EASY-Spray^TM^ electrospray source held at a constant temperature of 35*^◦^*C. Chromatography of peptides prior to mass spectral analysis was accomplished using a capillary emitter column (PepMap^○R^ C18, 3 *µ*M, 100 Å, 150 x 0.075 mm, Thermo Fisher Scientific) onto which 2 *µ*l of extracted peptides were automatically loaded. NanoHPLC system delivered solvents A: 0.1% (v/v) formic acid and B: 99.9% (v/v) ACN, 0.1% (v/v) formic acid were used at a flow rate of 0.5 *µ*l min^-1^ to load the peptides over a 30 minute period. A flow rate of 0.3 *µ*l min^-1^ was used to elute peptides directly into the nano-electrospray. A gradual gradient from 0% (v/v) B to 30% (v/v) B was used over 80 minutes and concluded with a 5 minute fast gradient from 30% (v/v) B to 50% (v/v) B at which time a 4 minute flash-out from 50-95% (v/v) B took place. A total run time of 150 minutes encompassed column conditioning at 95% B for 1 minute and equilibration at 100% A for 30 minutes. As peptides eluted from the HPLC-column/electrospray, source survey MS scans were acquired in the Orbitrap with a resolution of 120,000 folllwed by CID-type MS/MS with 2.0 AMU isolation and 10 ms activation time with 35% normalized collision energy fragmentation of 30 most intense peptides detected in the MS1 scan from 350 to 1800 m/z. Redundancy was limited by dynamic exclusion. Monoisotopic precursor selection and charge state screening were enabled and both +1 and undefined charge states were rejected.

#### 2.5.5 Protein Identification

Raw MS/MS data were converted to a.mgf file format using MSConvert (ProteoWizard: Open Source Software for Rapid Proteomic Tools Development) for downstream analysis. Resulting.mgf files were used to search against *Sphingomonasl sp.* LM7 Uniprot reference proteome database (UP000189383; 3,780 total entries, 09/23/2023 download) along with a list of common lab contaminants (172 total entries) using in-house *Mascot* search engine 2.7.0 (Matrix Science, Chicago, IL) with variable methionine oxidation, asparagine and glutamine deamidation plus fixed cysteine carbamidomethylation. Peptide mass tolerance was set at 10 ppm and fragment mass at 0.6 Da. Protein annotations, significance of identification, and spectral based quantification was done with the help of Scaffold software (version 4.11.0. Proteome Software Inc., Portland, OR). Peptide identifications were accepted if they could be established at greater than 99% probability to achieve an FDR less than 1% by the Scaffold Local FDR algorithm. Protein identifications were accepted if they could be established at FDR less then 1% and contained at least 2 identified peptides. Protein probabilities were assigned by the Protein Prophet algorithm. ^28^ Proteins that contained similar peptides and could not be differrentiated based on MS/MS analysis alone were grouped to satisfy the principles of parsimony. Proteins sharing significant peptide evidence were grouped into clusters.

### 2.6 Lectin Binding

A dot blot was conducted to check that the polysaccharide was still present after dialysis. 5 *µ*l of 50% w/v sample was pipetted onto a nitrocellulose membrane and allowed to dry for 15 minutes. The membrane was blocked with 25 ml of 5% bovine serum albumin (BSA) solution in TBST (100 ml 10X TBS buffer, 900 ml deionized water, 1 ml Tween-20). The membrane was then stained for 1 hour in 12 ml of 1 *µ*g mL^-1^ fluorescein conjugated Griffonia simplicifolia-II lectin solution with 1.25% BSA in TBST. Following the incubation, the membrane was rinsed twice with TBST solution for 10 minutes before imaging at a wavelength of 302 nm using an Azure 300 imager (Azure Biosystems, Dublin, CA).

### 2.7 Zeta Potential and Molecular Weight Measurements

Zeta potential and *M_W_* were measured using a Malvern Nano Zs Zetasizer (Malvern Pananalytical, Malvern, U.K.). Solutions were prepared in 2 ml tubes and given three hours to equilibrate before being transferred to polystyrene or zeta potential folded capillary cuvettes for measurement of either *M_W_* or zeta potential. Zeta potential was measured at a concentration of 0.1% with deionized water as the solvent. The zetasizer calculates the zeta potential by finding the electrophoretic mobility and then applying the Henry equation.^29^ The *M_W_* was measured using five consecutive concentrations, 0%, 0.025%, 0.05%, 0.075%, and 0.1%. Static light scattering was used to obtain a time-averaged intensity of scattered light. The *M_W_* was computed using the Rayleigh equation describing the relationship between the intensity of light scattered by a sample, its size, and *M_W_*.^30^

### 2.8 Fourier-Transform Infrared Spectroscopy

Fourier-Transform Infrared Spectroscopy (FTIR) measurements were performed on a Bruker Vertex FT-IR spectrometer (Bruker Corp., Billerica, MA). A spectral range of 600 cm^-1^ to 3500 cm^-1^ was selected. Approximately 5 mg of sample was loaded into the sample compartment to sufficiently cover the attenuated total reflectance (ATR) crystal. The sample chamber was vented with nitrogen before taking all reference and sample measurements.

### 2.9 Bicinchoninic acid (BCA) assay

A BCA protein assay was conducted to quantify protein content in the samples. A colorimetric kit was used measure the reduction of Cu^2+^ to Cu^1+^ by protein (Thermo Fisher Scientific). The color change was determined by measuring the optical density at a wavelength of 562 nm using a SpectraMax platereader (Danaher Life Sciences, Washington, DC).

### 2.10 Tensile Testing

Films were cast from 1% w/v promonan dissolved in deionized water and dried in ambient conditions. The dry sample mass was 1.7 ± 0.5 mg. The dry films had a density of 0.7 ± 0.1 g *cm^−^*^3^. Uniaxial tensile tests were conducted on a Zwick-Roell (Zwick-Roell, Germany) Z010 system with a 20 N capacity load cell. All tests were carried out using a strain rate of 0.05 *s^−^*^1^ with a pre-load of 0.1 N. Films were cast on a leveled surface and given 24 hours to dry at room temperature and humidity before being punched into dog bone shapes following ASTM D638 Type 5.

Digital image correlation (DIC) analysis was conducted by speckling tensile samples with white spray paint and back lighting the sample for imaging. Imaging was carried out using VIC-Snap-8 software and analyzed using VIC-2D Digital Image Correlation (Correlated Solutions, Irmo, SC).

### 2.11 Rheometry

Experiments were conducted using a TA Instruments DHR3 rheometer (Texas Instruments, Dallas, TA). A strain sweep at a frequency of 1 rad s^-1^ was used to determine the linear viscoelastic region and select a strain of 5% to be used for frequency sweeps. Both a strain sweep and a frequency sweep were conducted for each sample. A 40 mm parallel plate fixture was used with the gap set to 300 µm, requiring a sample volume of approximately 400 µl. The fixtures were thoroughly cleaned before use and between samples with deionized water and isopropyl alchohol and dried with compressed air. The sample was pipetted onto the bottom plate and the upper fixture lowered to the set gap value. A visual check confirmed that the amount of sample adequately filled the gap. Sweeps of the strain rate were used to produce the viscosity and stress graphs. Graphs presented are the average of three independent trials.

### 2.12 Differential Scanning Calorimetry (DSC)

Thermal analysis was conducted using a TA Instruments DSC Auto 2500 Differential Scanning Calorimeter. The sample was first weighed before being placed in a pan and sealed with a press. The temperature range of 90*^◦^*C to 400*^◦^*C was selected with a ramp rate of 10*^◦^*C min^-1^.

### 2.13 Thermogravimentric Analysis (TGA)

Thermogravimetric analysis was completed on a TA Instruments 5500 Thermogravimetric Analyzer (Texas Instruments, Dallas, TA). The sample was initially weighed and loaded using a platinum pan. The ramp rate was set to 10*^◦^*C min^-1^ and the gas supply was set to nitrogen.

### 2.14 Statistical Analysis

ANOVA tests were conducted to determine the probability of significant differences between groups. Two-tailed student t-tests were used to identify which groups were significant with a p-value of 0.01 to determine significance.

## 3 Results and Discussion

### 3.1 Pure Promonan Material Properties

Comparison between the chemical composition of promonan and other sphingans can provide insight into material properties including stiffness and thermal stability.^31^ Promonan is composed of a linear backbone of 3– and 4– linked glucose, and galactose and glucuronic acid residues with functional groups following the pattern seen for many sphingans and biopolysaccharides.^27^ The structure of promonan is similar to that of gellan gum, however, one of the main features of sphingans is the presence of rhamnose which is notably absent in promonan. The functional groups indicate high hydrogen bonding, a characteristic of carbohydrates. The carboxylic acid group gives the glucuronic acid a negative charge at pH 4.5 or higher and enables ionic cross-linking in the presence of positively charged bivalent ions.^32^ Despite the apparent similarity in chemical structures, analysis of the biosynthetic genes indicates that promonan has distinct biosynthesis pathways from other sphingans, such as gellan gum, and that the polymers represent distinct polysaccharide families.

Sphingans with similar chemical structure to promonan are used in healthcare applications due to their stiffness and biocompatibility. Gellan gum, with a Young’s modulus of 111 MPa, has been used as a stiffening component in antimicrobial films for wound healing. ^18,33^ We determined the suitability of promonan for similar applications through casting the material into a film and conducting uniaxial tensile testing as shown in Figure 2a and Supporting Video 1. For our material, promonan, the Young’s modulus is 282.06 ± 35.74 MPa while the fracture strength is 4.49 ± 1.28 MPa, with large variation due to sample defects initiating tearing, as shown in Supporting Information (SI) Figure 1. The high Young’s modulus for pure promonan films indicates that the material may be useful for providing mechanical structure in composite materials, such as in antimicrobial films for use in healthcare.

**Figure 2:**
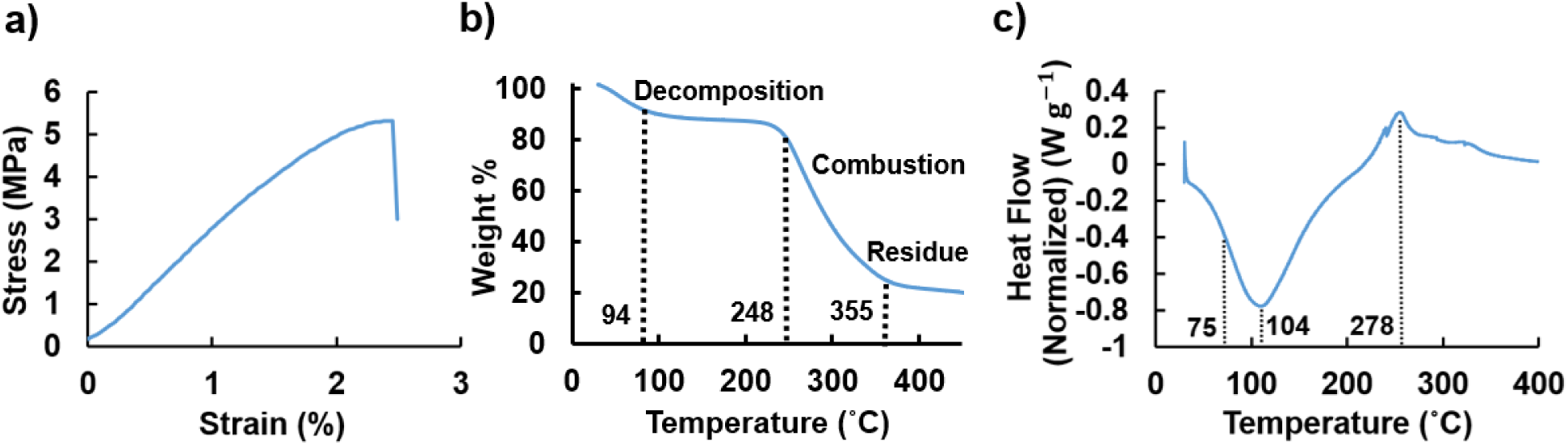
a) Representative stress-strain plot for tensile testing experiments with purified and cast promonan film. b) Thermal decomposition of the material shown through thermo-gravimetric analysis for purified polysaccharide. c) Analysis of the materials heat capacity tested using differential scanning calorimetry.

Sphingans have also been selected for many applications because of their thermal stability due to their ability to covalently cross-link, high *M_W_*, and stable glycosidic bonds.^34–36^ Both TGA and DSC were conducted to establish the thermal properties of this novel material and to enable comparison to fellow sphingans. The results of TGA show that weight loss occurs in two stages. There is an 11% decrease in weight in the first stage due to loss of absorbed and structural water as the temperature increases from 22.6*^◦^*C to 93.7*^◦^*C (Figure 2b).^37^ The main mass loss of 51% of the initial weight due to polysaccharide decomposition occurs between 248.1*^◦^*C and 355.3*^◦^*C. The sample weight gradually levels off with approximately 25% of the initial value remaining as ash. Therefore, the oxidation onset temperature is 248.1*^◦^*C and the maximum oxidation temperature is 355.3*^◦^*C. These results suggest that our material has good thermal stability, slightly greater than gellan gum. This may be due to the presence of exclusively beta glycosidic linkages in the proposed promonan structure which allows for a planar structure.^38,39^ DSC tests further support thermal stability and show consistency with the TGA data. Figure 2c shows an endothermic event at 104.4*^◦^*C possibly due to water evaporation, which would correspond to the TGA results. The DSC tests also identify the glass transition temperature as 75.1*^◦^*C, crystallization as occurring between 104.4*^◦^*C and 277.5*^◦^*C and a melting temperature of 277.5*^◦^*C. These results correspond well with the second mass loss event in TGA with the exothermic peak of 277.5*^◦^*C as seen in the Figure 2c. These temperatures follow the patterns for other thermally stable biologically produced polysaccharides for both TGA and DSC.^40^

### 3.2 Material Properties of Hydrated Promonan

Bacterial polysaccharides are widely used in a hydrated state. Structural characteristics including high *M_W_* and a negative surface charge provide emulsion stability for many applications.^41,42^ The *M_W_* and zeta potential of promonan were analyzed to gain insight into the properties of the material in a hydrated state. The *M_W_* of promonan, 579 ± 120 kDa is similar to gellan gum, which has an *M_W_* of approximately 500 kDa.^41^ High *M_W_* typically correlates with chemical resistance as it takes more main chain scission events before the material strength is affected. ^43,44^ Lastly, the high *M_W_* indicates a higher viscosity at room temperature for water based solutions at a given w/v ratio, as longer chains resist flow due to higher entanglement. The zeta potential is –31.8 ± 2.7 mV, similar to other sphingans such as gellan gum, which has a zeta potential of –29.1 ± 3.0 mV.^42^ A negative zeta potential value is expected due to the carboxylic acid groups. The zeta potential indicates the potential stability of the colloidal system. Typically, a value below –30 mV or above +30 mV is considered stable, whereas materials with zeta potentials within this range tend to aggregate within a solution. A zeta potential value below –30 mV for promonan therefore, indicates that promonan particles are likely to repel each other rather than aggregating and flocculating. This is ideal for many applications where aggregation would not be desirable and demonstrates that the material dispersion is stable in a hydrated state, expanding the possible uses of promonan.

Bacterial polysaccharides are often used as thickening agents where large changes in viscosity due to small increases in the w/v ratio are of interest. The experiments show that promonan acts as a gel at w/v ratios over 1%. Figure 3a shows the effect of varying water content on the dynamic moduli of promonan measured using rheology. A 1% w/v promonan gel can be conveniently pipetted and handled for testing while a 3% w/v gel is too viscous to be pipetted. In contrast, the 0.5% ratio does not gelate and maintains solution like behavior throughout the frequency sweep. A small increase from 0.5% to 3% w/v ratio changes the storage modulus by several orders of magnitude.

**Figure 3:**
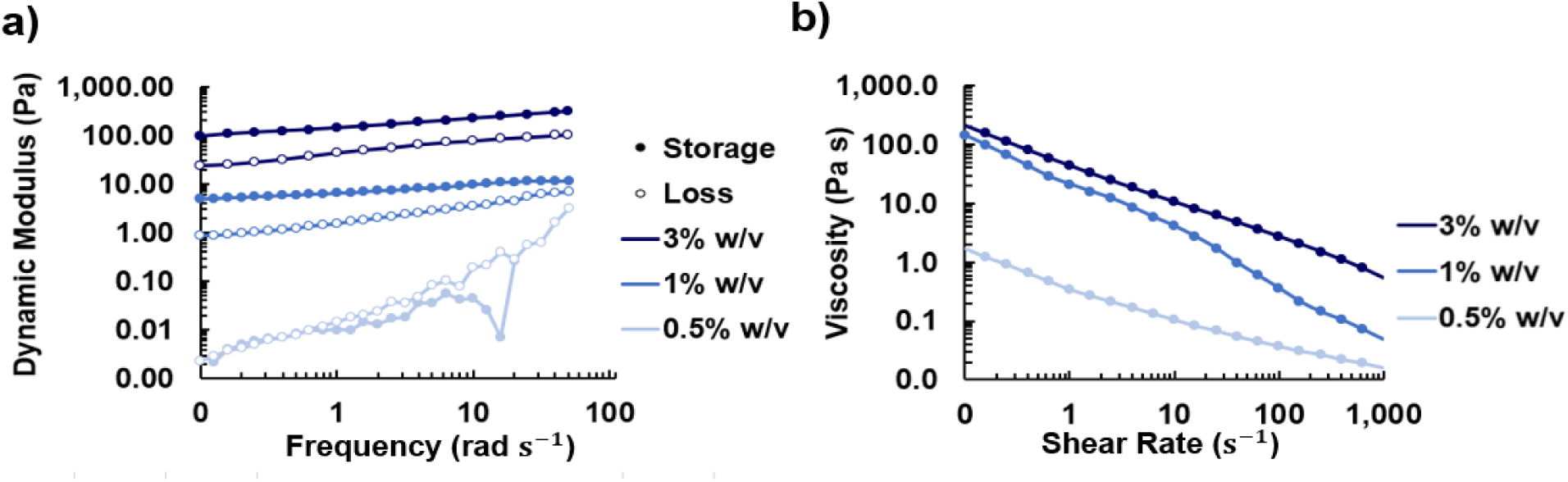
a) Increasing the w/v percentage of promonan hydrated in deionized water increases the dynamic moduli of the gel by several orders of magnitude and shifts the material state from solution-like to gel-like at higher values b) Increasing the w/v percentage of promonan increases the viscosity, however at all concentrations the gel is shear thinning.

The changes in viscosity as water content is altered can be compared to study the changes in the material behavior. These properties are of particular interest for polysaccharides used as stabilizers, binders and preservatives in the food industry.^11,14,16^ A flow sweep was conducted by varying the shear rate, as shown in Figure 3b for the 0.5%, 1% and 3% w/v fractions. The figure indicates shear thinning viscosity for all three concentrations. The negative surface charge causes molecules to repel each other, inhibiting flow and increasing the viscosity at low shear rates. Promonan would make an effective thickening agent and could be used in food-processing, bioprinting, and cosmetic applications.

### 3.3 Tailoring Rheological Properties with Ions

In many applications it is desirable to be able to tune properties such as material stiffness. One prominent example is bioprinting, which requires a shear thinning material and a stiffness of approximately 0.5 kPa.^45–47^ Previous studies have explored bioprinting of various polysaccharides, like alginate, cross-linked with ions, such as calcium, for tissue culturing scaffolds due to their biocompatibility and rheological properties. ^45,48^ We demonstrate in this study that the dynamic moduli of promonan increases with the addition of ions by adding a salt to hydrated samples to be comparable to gels used for bioprinting.^48^ Bivalent calcium ions cross-link carboxylic acid sites resulting in an increase to both the storage and loss moduli by approximately one order of magnitude (Figure 4a). Furthermore, we show that Ethylenediamine tetraacetic acid (EDTA) can be used to chelate ions and decrease the dynamic moduli, a desirable functionality for tuning a bioink to be a substrate for a particular cell type. As expected, the monovalent sodium ions, which should not enable ionic cross-linking, do not cause a statistically significant increase to the dynamic moduli (Figure 4b).

**Figure 4:**
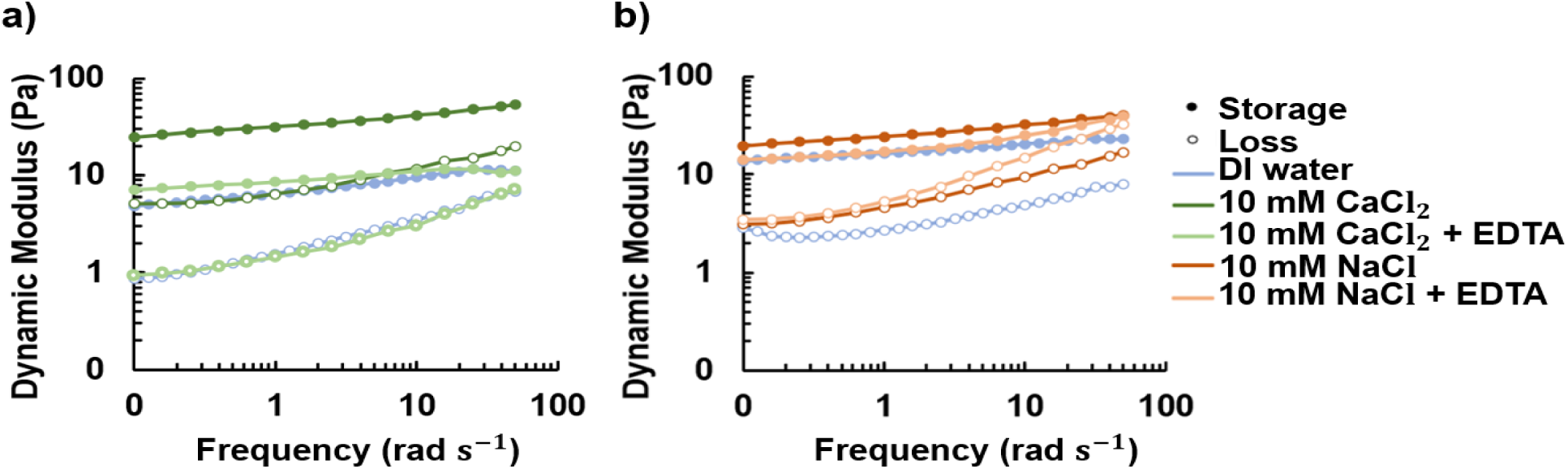
a) The addition of bivalent ions results in a significant increase in the dynamic moduli while the addition of EDTA chelates the ions (p-value *<* 0.01), reducing the dynamic moduli. b) The addition of monovalent ions results in a small increase in dynamic moduli that cannot be reversed with EDTA, which does not chelate monovalent ions. These changes were not statistically significant (p-value *>* 0.01).

### 3.4 The Impact of the Purification Process on Material Properties

Biomanufactured materials typically undergo many purification steps. However, it is unclear how the purification process alters the material properties and whether the process could be shortened while still yielding a usable material. We investigated methods of shortening the purification process to accelerate production, increase the material yield and facilitate potential future manufacturing of the material. The first method follows the bacteria culturing step with dialysis to remove undesired low *M_W_* proteins and culture media components and eliminate the need for enzymatic purification steps (Figure 5a). The material yield, shown in Figure 5b, drops significantly when salts and sucrose from the growth media are removed via dialysis with an MWCO of 3.5 kDa. Dialysis significantly decreased the material yield through removing culture media components. Although the value of 100 mg L^-1^ for the final purified material yield is higher than for the 100 kDa MWCO dialyzed sample with a yield of 77 mg L^-1^, the values were not significantly different. A dialysis MWCO of 50 kDa or higher greatly reduces the protein content by over 15%, as shown in Figure 5c. A graph without the unpurified sample is included in Figure S4a. A dot blot was also performed to demonstrate that the polysaccharide is still present following dialysis for each MWCO (Figure S4b). Note that the sample dialyzed at 100 kDa was too viscous to permeate into the nitrocellulose membrane.

**Figure 5:**
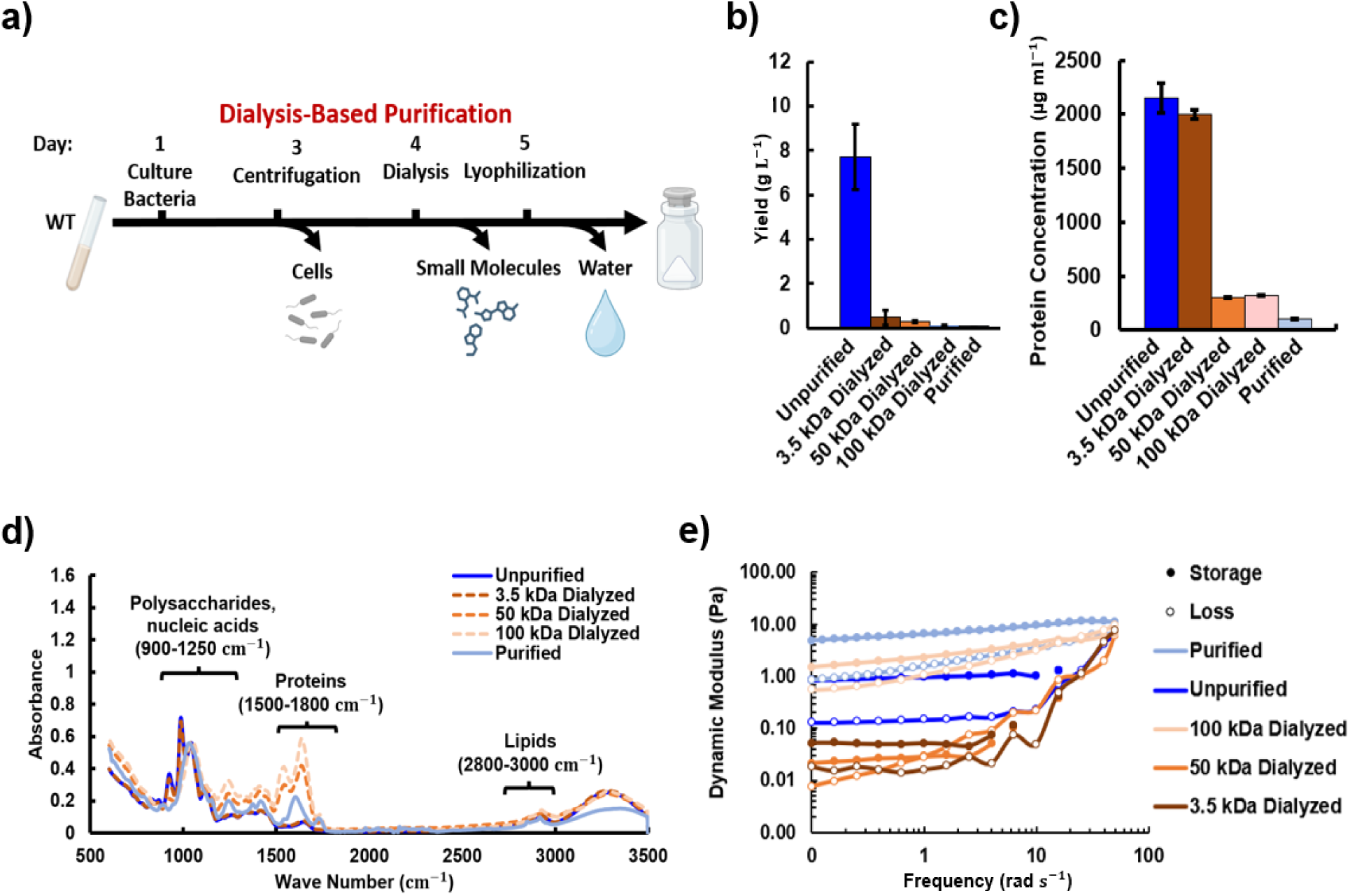
a) The dialysis-based purification process consists of three steps; centrifugation, dialysis and lyophilization to remove cells, small molecules and water respectively. b) The reduction in sample yield as material is lost at various points in purification and with different MWCO values during dialysis. c) The protein content within the samples decreases with the dialysis levels with the purified sample having the lowest quantity. d) FTIR comparison of samples with varying levels of purification normalized to the C-O peak at 1050 cm^-1^. e) Comparison of the rheological behaviour between unpurified, purified and dialyzed samples.

The changes in composition were further analyzed via FTIR for each of the dialyzed, unpurified and purified samples (Figure 5d). Regions for lipids, proteins and polysaccharides are labeled based on literature on unpurified biofilms. ^49^ The spectra show a gradual shift from the unpurified spectra toward the purified spectra, indicating removal of proteins and other substances. The typical spectral pattern for polysaccharides^50^ is apparent in all cases. The unpurified sample closely matches peaks of the known spectra for sucrose, which is not surprising given the high amount of sucrose used in the growth media for *Sphingomonas sp.* LM7.^51^ The peaks in Figure 5d indicate that the functional groups found include both primary and secondary alcohols (separate peaks) at 3350 cm^-1^ and 2918 cm^-1^, amine salts and strong O-H, C-H, C-O stretches in the spectra (1031 cm^-1^). The peak at approximately 3200 cm^-1^, likely from water absorption, remains small indicating that there is likely little effect on O-H bond stretching.

Rheology was used measure how compositional changes altered stiffness. Figure 5e shows that the rheological stiffness of the unpurified material is significantly lower than the purified material, but does remain gel-like with the storage modulus above the loss modulus throughout the frequency sweep. The presence of proteins, sugars, deoxyribonucleic acid (DNA) and ribonucleic acid (RNA) with lower *M_W_* values than promonan likely results in a lower storage and loss modulus for the unpurified sample. However, the components of the material may also interact to increase material stiffness. Sucrose, for example, has been shown to stabilize proteins and increase the viscosity of protein solutions through the concept of preferential exclusion and hydration. ^52^ The sugars are excluded from the volume around the protein leading to preferential hydration of the protein shell region. This results in a more compact and stable protein folding state and increases the viscosity. Protein in the unpurified sample accounts for of 21.5% of the solid material. In contrast, the protein concentration in the purified material is estimated to be 1%. The protein concentrations for the dialyzed samples with MWCOs of 3.5 kDa, 50 kDa and 100 kDa are 20%, 3% and 3% respectively. Figure 5e shows a further decrease in moduli and transition to fluid-like behavior when the large amount of sucrose is taken out with dialysis at 3.5 kDa MWCO. The dynamic moduli are found to increase relative to the 3.5 kDa MWCO material when a higher MWCO for dialysis is used. Figure 5e shows a transition back to gel-like behavior when 50 kDa is used. The values for the storage and loss moduli are largely recovered, relative to the enzymatically purified material, when using the 100 kDa MWCO. These results indicate that mechanical properties close to the enzymatically purified material could be obtained using a shorter, dialysis-based purification process, especially if the ability to further increase dynamic moduli with divalent ions or decreasing water content is taken into account.

### 3.5 Purification via Ultracentrifugation

The presence of proteins, which have a lower *M_W_* than promonan, may be contributing to the decrease in dynamic moduli of the dialyzed material compared to the enzymatically-purified material. To understand which proteins might alter the material stiffness, we analyzed proteins present in the spent medium of LM7 cultures by performing mass spectrometry-based identification of bands excised from an SDS-PAGE gel (Figure 6a). Many of the proteins identified in the analysis were predicted to be located in the gram-negative outer membrane, suggesting that they were released from the cell in outer membrane vessicles (OMVs). The most abundant proteins identified were three different TonB-dependent receptor proteins, an OmpA family protein, Flagellin, outer membrane (OM) lipoprotein pal and a 17 kDa surface antigen. We therefore explored ultracentrifugation as a means of removing OMVs to lower the amount of protein in the sample. Although ultracentrifugation did not significantly alter the material yield compared to the unpurified material, it was found to successfully decrease the protein content 6b, c. Coupling ultracentrifugation with dialysis produced a yield of 0.16 g L^-1^ compared with 0.27 g L^-1^ for 100 kDa dialysis alone. This combined ultracentrifugation and 100 kDa dialysis also resulted in a further decrease in protein abundance relative to the amount of polysaccharide of interest as seen in 6d. The protein to polysaccharide ratio for the ultracentrifuged 100 kDa dialyzed sample was 0.66, slightly lower than that of the 100 kDa dialyzed sample at 0.70.

**Figure 6:**
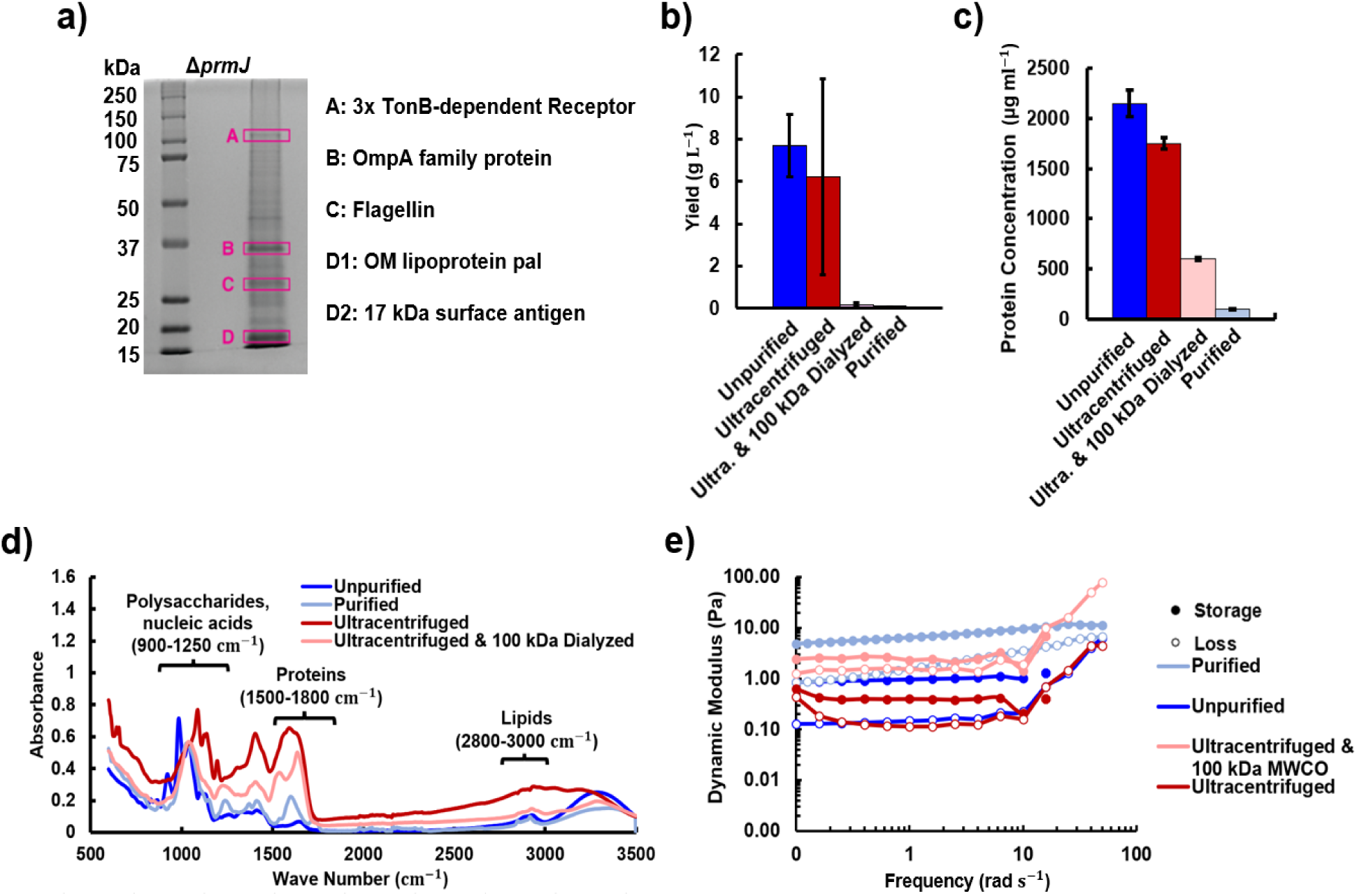
a) Analysis of secreted proteins for *Sphingomonas sp.* LM7. b) Sample yield for unpurified, ultracentrifuged, ultracentrifuged and dialyzed, and enzymatically purified material. c) Protein quantification for the unpurified, ultracentrifuged, and enzymatically purified samples. d) FTIR comparison of samples undergoing ultracentrifugation normalized to the C-O peak at 1050 cm^-1^. e) Comparison of the rheological behaviour between samples showing an increase in the dynamic moduli when unpurified material undergoes ultracentrifugation and a further increase with subsequent dialysis.

The changes in material composition due to ultracentrifugation alter the rheological properties as seen in 6e. Results of the frequency sweep show a slight decrease in the moduli between the unpurified material and the ultracentrifuged material which could be due to the high shear stresses applied during centrifugation (Figure 6e).^53^ High shear stresses can decrease the *M_W_* and viscosity of polymer solutions which may decrease the dynamic moduli. The shear stresses may also facilitate subsequent dialysis by shearing polymer chains. Lectin binding demonstrates that the polysaccharide is not removed by ultracentrifugation (Figure S5a). Figure 6e shows that when ultracentrifugation is combined with dialysis, the material moduli improve significantly while yield significantly decreases, likely due to the removal of protein, lipids and DNA as shown in Figures 6c, d and Figure S7.

### 3.6 Protein Knockout Purification Pathway

One advantage to producing materials in genetically tractable organisms is the potential to make mutations that simplify the purification process. We explored the potential of this strategy by deleting the gene for a protein that is secreted in high abundance by LM7 based on the identified secreted proteins shown in Figure 6a. We chose to delete the gene for Δ*fliC*, as this was the only protein identified in the supernatant that was not predicted to be contained in OMVs. A Δ*fliC* strain was developed that still produced promonan, but not *fliC*, the protein that polymerizes to form filaments for bacterial flagella. Analysis of the overall material yield and protein content indicate a decrease in the total material and protein abundance (Figures 7b, c). The purified *fliC* sample yield, 0.02 *µ*g L^-1^ was approximately one fifth that of the wild type purified material. Interestingly, in this case, dialysis did not further decrease the protein content but is shown to decrease yield through the removal of solute from the growth media. Further study of which proteins are produced by the strain, as shown in Figure S6a, and the resulting properties may lend insight into what additional protein knockouts or modifications could be used to alter the material composition. FTIR, shown in Figure 7d, indicates that the protein to polysaccharide ratio is much greater for both the Δ*fliC* and the Δ*fliC* dialyzed samples than the enzymatically purified material. The lipid to polysaccharide ratio for both the Δ*fliC* and the Δ*fliC* dialyzed samples is also higher than both the purified and unpurified samples however, the intensities of the lipid peaks remain low relative to the protein and polysaccharide peaks.

**Figure 7:**
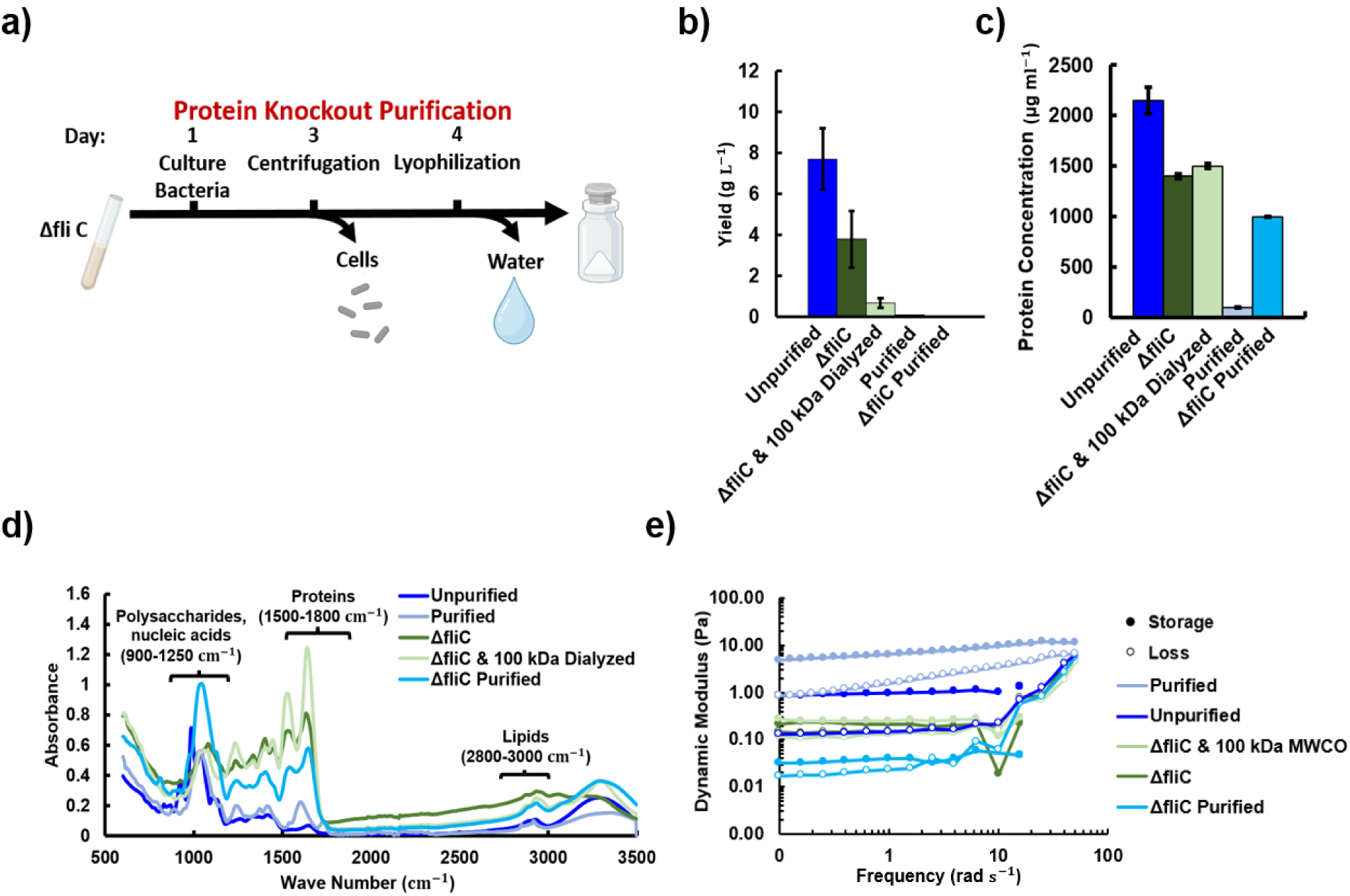
a) The protein knockout purification pathway consists of culturing the Δ*fliC* strain before removing cells and water through centrifugation and lyophilization. b) Sample yield for unpurified, Δ*fliC*, Δ*fliC* dialyzed, enzymatically purified and Δ*fliC* purified material. c) Protein quantification for the unpurified, Δ*fliC*, and Δ*fliC* dialyzed, purified and Δ*fliC* purified samples. d) FTIR comparison of samples from wild type and Δ*fliC* strains normalized to the C-O peak at 1050 cm^-1^. e) Comparison of the rheological behaviour between samples showing little change in the dynamic moduli for material produced from the wild type and Δ*fliC* strains.

The changes in composition between the wild type and Δ*fliC* produced material offer insight into the different rheological behaviors observed. Figure 7e shows that the dynamic moduli of the protein knockout material are decreased compared to the unpurified material, possibly due to the increased protein to polysaccharide ratio and lipid to polysaccharide ratio or the production of the signalling molecule cyclic dimeric guanosine monophosphate (c-di-GMP). Mutating genes that affect the flagellum have been shown to influence mechanosensing pathways and activate the production of c-di-GMP,^54,55^ which can alter enzymes related to biofilm production.^55^ The purified Δ*fliC* material also had decreased dynamic moduli compared to the purified material despite having similar protein to polysaccharide ratios. This may point to the role of signalling molecules in mechanics however, the Δ*fliC* mutant may also be secreting a less soluble form of promonan as has been seen with other mutants of the LM7 strain.^27^ Dialyzing the spent media from the Δ*fliC* strain removes the viscosity enhancing effect of sucrose but also eliminates low molecular weight species, such as charged ions, resulting in little overall change in stiffness. Charged ions are thought to alter protein-protein interactions and ultimately decrease solution viscosity. ^52^ The 100 kDa MWCO dialyzed wild type strain had dynamic moduli that were approximately two orders of magnitude stiffer than the 100 kDa dialyzed Δ*fliC* strain. These results demonstrate that genetic modification can be applied to alter the material properties and reduce the required purification steps. However, additional research is needed to identify genetic modifications capable of increasing the rheological stiffness.

## 4 Conclusions

In this paper we have successfully characterized a novel bacterial polysaccharide, providing an overview of thermal, rheological and mechanical properties. It is interesting that promonan and sphingans, such as gellan gum, display similar material parameters. This suggests that polysaccharides with divergent biosynthesis pathways can produce polymers with similar properties. The potential applications of the material for films and bioinks was demonstrated via tensile testing of films and rheological tests of hydrogels. Salt was added to hydrated promonan samples to investigate the effectiveness of ions in crosslinking the gel to tailor the stiffness. This study demonstrated that the lengthy purification process can be shortened by over 50% while still yielding gel state polymers at 1% w/v ratio. Alternative purification processes using dialysis and genetic modification were compared to improve the purification time and user-friendliness. Purification via dialysis with a MWCO of 100 kDa provided the closest rheological properties to those of the enzymatically purified material. This paper provides an overview of material properties, identifies advantages such as tailorable properties for applications and highlights the material’s manufacturability through various processing methods.

Our work identified material properties that might be suitable for an array of applications in healthcare, biomanufacturing and food processing. Future research is required to demonstrate these applications as well as investigate further material modifications, such as functionalization for bio-remediation or as an antimicrobial film for use in environmental or healthcare applications.

## Acknowledgement

This work was supported through the NSERC CGS-D scholarship (E. W. v. W.), the Beckman Young Investigator award (D. M. H.) and NIH R35GM150652 (D. M. H). This work was supported and performed in part at the Engineered Living Materials institute. This work was performed in part at the Cornell NanoScale Facility, a member of the National Nanotechnology Coordinated Infrastructure (NNCI), which is supported by the National Science Foundation (Grant NNCI69 2025233). The authors acknowledge the use of facilities and instrumentation supported by NSF through the Cornell University Materials Research Science and Engineering Center DMR-1719875.

## Supporting Information Available

The following files are available free of charge.

Supporting information.pdf: Supplementary figures S1-S8 including uniaxial tensile tests, strain sweeps for all rheological tests, lectin binding assay results, DNA content analyses. Supplementary tables listing strains and plasmids used in this study.

Supplementary Video.mp4: Video of digital image correlated tensile test.

## TOC Graphic

**Figure 8:**
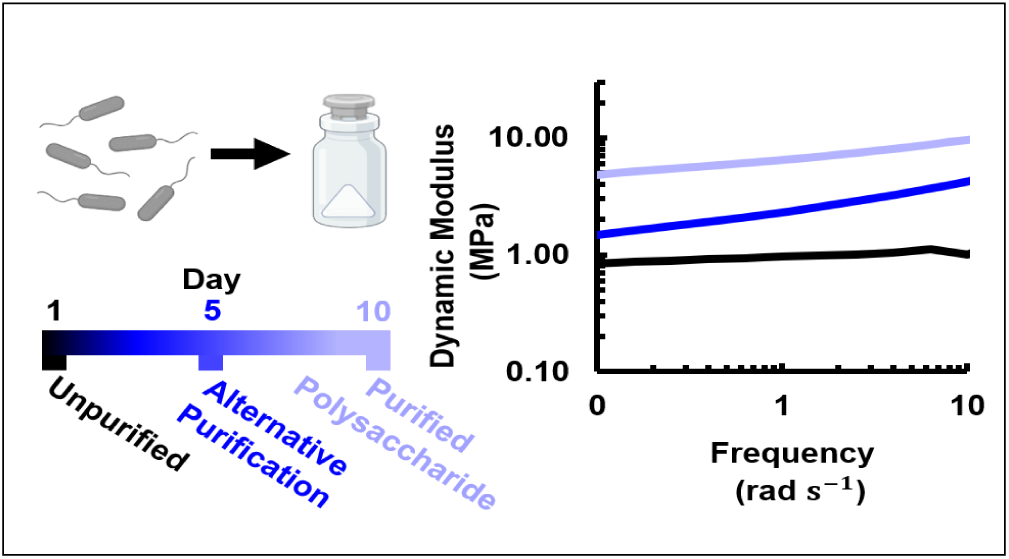
Changing the material composition through alternative processing methods reduces production time and enables tuning of material stiffness for extracellular polymeric substances isolated from *Sphingomonas sp.* LM7.

## Notes

### Competing Interest Statement

The authors have declared no competing interest.

## References

(1) The White House Office of Science {and} Technology Policy, Bold Goals for U.S. Biotechnology and Biomanufacturing: Harnessing Research and Development to Further Societal Goals.

(2) Meng, F.; Ellis, T. The second decade of synthetic biology: 2010–2020. 11, 5174.

(3) Wang, W.; Zheng, G.; Lu, Y. Recent Advances in Strategies for the Cloning of Natural Product Biosynthetic Gene Clusters. 9.

(4) Zhang, Z.; Parker, M.; Liao, K.; Cao, J.; Waghmare, A.; Breda, J.; Matsumura, C.; Eley, S.; Roumeli, E.; Patel, S.; Iyer, V. Biodegradable Interactive Materials. http://arxiv.org/abs/2404.03130.

(5) Nguyen, P. Q.; Courchesne, N.-M. D.; Duraj-Thatte, A.; Praveschotinunt, P.; Joshi, N. S. Engineered Living Materials: Prospects and Challenges for Using Biological Systems to Direct the Assembly of Smart Materials. 30, 1704847.

(6) Rehm, B. H. A. Bacterial polymers: biosynthesis, modifications and applications. 8, 578–592.

(7) Limoli, D. H.; Jones, C. J.; Wozniak, D. J. Bacterial Extracellular Polysaccharides in Biofilm Formation and Function. 3, 10.1128/microbiolspec.MB-0011-2014.

(8) Palaniraj, A.; Jayaraman, V. Production, recovery and applications of xanthan gum by Xanthomonas campestris. 106, 1–12.

(9) Singh, S. et al. Carbohydrates from Pseudomonas aeruginosa biofilms interact with immune C-type lectins and interfere with their receptor function. 7, 1–14.

(10) Kaur, V.; Bera, M. B.; Panesar, P. S.; Kumar, H.; Kennedy, J. F. Welan gum: Microbial production, characterization, and applications. 65, 454–461.

(11) McGuffey, J. C.; Leon, D.; Dhanji, E. Z.; Mishler, D. M.; Barrick, J. E. Bacterial Production of Gellan Gum as a Do-It-Yourself Alternative to Agar. 19, 19.2.74.

(12) Wahid, F.; Zhao, X.-J.; Zhao, X.-Q.; Ma, X.-F.; Xue, N.; Liu, X.-Z.; Wang, F.-P.; Jia, S.-R.; Zhong, C. Fabrication of Bacterial Cellulose-Based Dressings for Promoting Infected Wound Healing. 13, 32716–32728.

(13) Brar, V.; Kaur, G. Preparation and Characterization of Polyelectrolyte Complexes of Hibiscus esculentus (Okra) Gum and Chitosan. 2018, 4856287.

(14) Moradali, M. F.; Rehm, B. H. A. Bacterial biopolymers: from pathogenesis to advanced materials. 18, 195–210.

(15) Li, Y.; Xu, L.; Gong, H.; Ding, B.; Dong, M.; Li, Y. A Microbial Exopolysaccharide Produced by Sphingomonas Species for Enhanced Heavy Oil Recovery at High Temperature and High Salinity. 31, 3960–3969, Publisher: American Chemical Society.

(16) Huang, H.; Lin, J.; Wang, W.; Li, S. Biopolymers Produced by Sphingomonas Strains and Their Potential Applications in Petroleum Production. 14, 1920.

(17) Seymour, R. B.; Carraher, C. E. In Structure—Property Relationships in Polymers; Seymour, R. B., Carraher, C. E., Eds.; Springer US, pp 83–93.

(18) Freitas, F.; Alves, V. D.; Reis, M. A. M. Advances in bacterial exopolysaccharides: from production to biotechnological applications. 29, 388–398.

(19) Li, H.; Jiao, X.; Sun, Y.; Sun, S.; Feng, Z.; Zhou, W.; Zhu, H. The preparation and characterization of a novel sphingan WL from marine Sphingomonas sp. WG. 6, 37899.

(20) Shi, L. Bioactivities, isolation and purification methods of polysaccharides from natural products: A review. 92, 37–48.

(21) Stouten, G. R.; Hogendoorn, C.; Douwenga, S.; Kilias, E. S.; Muyzer, G.; Kleerebezem, R. Temperature as competitive strategy determining factor in pulse-fed aerobic bioreactors. 13, 3112.

(22) Seviour, R. J.; McNeil, B.; Fazenda, M. L.; Harvey, L. M. Operating bioreactors for microbial exopolysaccharide production. 31, 170–185.

(23) Singhania, R. R.; Patel, A. K.; Tsai, M.-L.; Chen, C.-W.; Di Dong, C. Genetic modification for enhancing bacterial cellulose production and its applications. 12, 6793–6807.

(24) Jennings, L. K.; Storek, K. M.; Ledvina, H. E.; Coulon, C.; Marmont, L. S.; Sadovskaya, I.; Secor, P. R.; Tseng, B. S.; Scian, M.; Filloux, A.; Wozniak, D. J.; Howell, P. L.; Parsek, M. R. Pel is a cationic exopolysaccharide that cross-links extracellular DNA in the Pseudomonas aeruginosa biofilm matrix. 112, 11353–11358.

(25) Xu, L.; Xu, G.; Liu, T.; Chen, Y.; Gong, H. The comparison of rheological properties of aqueous welan gum and xanthan gum solutions. 92, 516–522.

(26) Miyoshi, E.; Takaya, T.; Nishinari, K. Gel-sol transition in gellan gum solutions. I. Rheological studies on the effects of salts. 8, 505–527.

(27) Goetsch, A. G.; Ufearo, D.; Keiser, G.; Heiss, C.; Azadi, P.; Hershey, D. M. A novel exopolysaccharide pathway from a freshwater Sphingomonas isolate. https://www.biorxiv.org/content/10.1101/2023.11.03.565537v1.

(28) Nesvizhskii, A. I.; Keller, A.; Kolker, E.; Aebersold, R. A Statistical Model for Identifying Proteins by Tandem Mass Spectrometry. 75, 4646–4658.

(29) Kaszuba, M.; Corbett, J.; Watson, F. M.; Jones, A. High-concentration zeta potential measurements using light-scattering techniques. 368, 4439–4451.

(30) Kruppa, B.; Strube, G.; Gerlach, C. In Optical Measurements: Techniques and Applications; Mayinger, F., Feldmann, O., Eds.; Heat and Mass Transfer; Springer, pp 99–116.

(31) West, T. P. Synthesis of the Microbial Polysaccharide Gellan from Dairy and Plant-Based Processing Coproducts. 2, 234–244.

(32) Nakashima, I.; Kishida, A.; Takaoka, Y.; Morisada, S.; Ohto, K.; Kawakita, H.; Iwasaki, W.; Sathuluri, R. R.; Miyazaki, M. Adsorption and Elution of Glucuronic Acid and Chondroitin Sulfate Using Amino-Group-Containing Spherical Gel. 63, 69–75.

(33) Mohd Azam, N. A. N.; Amin, K. A. M. The Physical and Mechanical Properties of Gellan Gum Films Incorporated Manuka Honey as Wound Dressing Materials. 209, 012027.

(34) Xu, Z.; Li, Z.; Jiang, S.; Bratlie, K. M. Chemically Modified Gellan Gum Hydrogels with Tunable Properties for Use as Tissue Engineering Scaffolds. 3, 6998–7007.

(35) Lu, X. Changes in the structure of polysaccharides under different extraction methods. 4, e82.

(36) Zhang, J.; Huo, H.; Zhang, L.; Yang, Y.; Li, H.; Ren, Y.; Zhang, Z. Effect of High-Temperature Hydrothermal Treatment on the Cellulose Derived from the Buxus Plant. 14, 2053.

(37) Vendruscolo, C. W.; Ferrero, C.; Pineda, E. A. G.; Silveira, J. L. M.; Freitas, R. A.; Jiménez-Castellanos, M. R.; Bresolin, T. M. B. Physicochemical and mechanical characterization of galactomannan from Mimosa scabrella: Effect of drying method. 76, 86–93.

(38) Sukumar, S.; Arockiasamy, S.; Chemmattu Moothona, M. Optimization of cultural conditions of gellan gum production from recombinant Sphingomonas paucimobilis ATCC 31461 and its characterization. 9.

(39) Nep, E. I.; Conway, B. R. Characterization of Grewia Gum, a potential pharmaceutical excipient. *1*, 30–40.

(40) Lin, C.-P.; Tsai, S.-Y. Differences in the Moisture Capacity and Thermal Stability of Tremella fuciformis Polysaccharides Obtained by Various Drying Processes. 24, 2856.

(41) Feketshane, Z.; Alven, S.; Aderibigbe, B. A. Gellan Gum in Wound Dressing Scaffolds. 14, 4098.

(42) Duarte, L. G.; Alencar, W. M.; Iacuzio, R.; Silva, N. C.; Picone, C. S. Synthesis, characterization and application of antibacterial lactoferrin nanoparticles. 5, 642–652.

(43) Vaganov, G.; Simonova, M.; Romasheva, M.; Didenko, A.; Popova, E.; Ivan’kova, E.; Kamalov, A.; Elokhovskiy, V.; Vaganov, V.; Filippov, A.; Yudin, V. Influence of Molecular Weight on Thermal and Mechanical Properties of Carbon-Fiber-Reinforced Plastics Based on Thermoplastic Partially Crystalline Polyimide. 15, 2922.

(44) Levy, A.; Wang, F.; Lang, A.; Galant, O.; Diesendruck, C. E. Intramolecular Cross-Linking: Addressing Mechanochemistry with a Bioinspired Approach. 56, 6431–6434.

(45) GhavamiNejad, A.; Ashammakhi, N.; Wu, X. Y.; Khademhosseini, A. Crosslinking Strategies for Three-Dimensional Bioprinting of Polymeric Hydrogels. 16, e2002931.

(46) Sharma, R.; Kirsch, R.; Valente, K. P.; Perez, M. R.; Willerth, S. M. Physical and Mechanical Characterization of Fibrin-Based Bioprinted Constructs Containing Drug-Releasing Microspheres for Neural Tissue Engineering Applications. 9, 1205.

(47) Bay, R. K.; Hancock, A. M.; Dill-Macky, A. S.; Luu, H. N.; Datta, S. S. Author Spotlight: Studying Bacterial Growth in 3D Hydrogel Matrices. e66166.

(48) Gorroñogoitia, I.; Urtaza, U.; Zubiarrain-Laserna, A.; Alonso-Varona, A.; Zaldua, A. M. A Study of the Printability of Alginate-Based Bioinks by 3D Bioprinting for Articular Cartilage Tissue Engineering. 14, 354.

(49) Di Martino, P. Extracellular polymeric substances, a key element in understanding biofilm phenotype. 4, 274–288.

(50) Gómez-Ordóñez, E.; Rupérez, P. FTIR-ATR spectroscopy as a tool for polysaccharide identification in edible brown and red seaweeds. 25, 1514–1520.

(51) Patil, N. V.; Netravali, A. N. Multifunctional sucrose acid as a ‘green’ crosslinker for wrinkle-free cotton fabrics. 27, 5407–5420.

(52) He, F.; Woods, C. E.; Litowski, J. R.; Roschen, L. A.; Gadgil, H. S.; Razinkov, V. I.; Kerwin, B. A. Effect of Sugar Molecules on the Viscosity of High Concentration Monoclonal Antibody Solutions. 28, 1552–1560.

(53) Ram, A.; Kadim, A. Shear degradation of polymer solutions. 14, 2145–2156.

(54) Zorraquino, V.; García, B.; Latasa, C.; Echeverz, M.; Toledo-Arana, A.; Valle, J.; Lasa, I.; Solano, C. Coordinated Cyclic-Di-GMP Repression of Salmonella Motility through YcgR and Cellulose. 195, 417–428.

(55) Wu, D. C.; Zamorano-Sánchez, D.; Pagliai, F. A.; Park, J. H.; Floyd, K. A.; Lee, C. K.; Kitts, G.; Rose, C. B.; Bilotta, E. M.; Wong, G. C. L.; Yildiz, F. H. Reciprocal c-diGMP signaling: Incomplete flagellum biogenesis triggers c-di-GMP signaling pathways that promote biofilm formation. 16, e1008703.

