## Supporting Information for "Engineering Bacterial Biomanufacturing: Characterization and Manipulation of *Sphingomonas sp.* LM7 Extracellular Polymers"

### Supporting Information: Engineering Bacterial Polymers for Biomanufacturing: Characterization and Manipulation of *Sphingomonas sp. LM7* Extracellular Polysaccharide

Ellen W. van Wijngaarden †, Alexandra G. Goetsch ‡, Ilana L. Brito ¶, David M. Hershey ‡, and Meredith N. Silberstein †

†*Sibley School of Mechanical and Aerospace Engineering, Cornell University, Ithaca, NY*
‡*Department of Bacteriology, University of Wisconsin-Madison, Madison, WI*
¶*Meinig School of Biomedical Engineering, Cornell University, Ithaca, NY 14853, USA*


Phone: 607/255-5063

### 1 Supporting Figures

#### 1.1 Uniaxial Tensile Tests

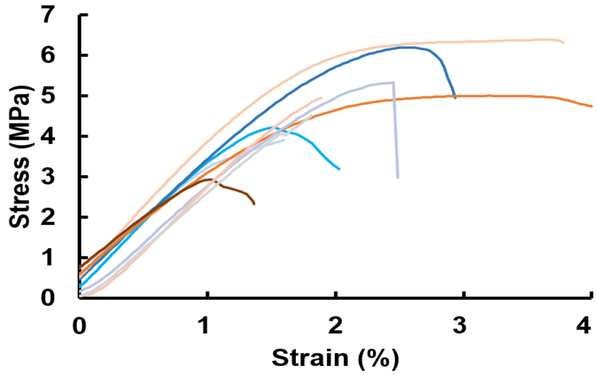

Figure S1: Tensile testing trials for films showing variation in the fracture strength of the material. Young's Modulus is consistent among tests.

#### 1.2 The Effect of Water Content on Rheological Properties: Strain Sweep

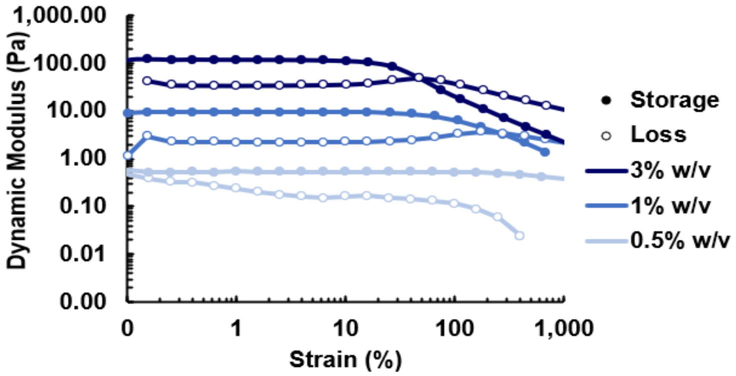

Figure S2: Strain sweep results comparing sample water content showing the gel-like behavior. Dynamic moduli increase for higher w/v ratios.

#### 1.3 How Bivalent and Monovalent Ions alter Rheological Properties: Strain Sweep

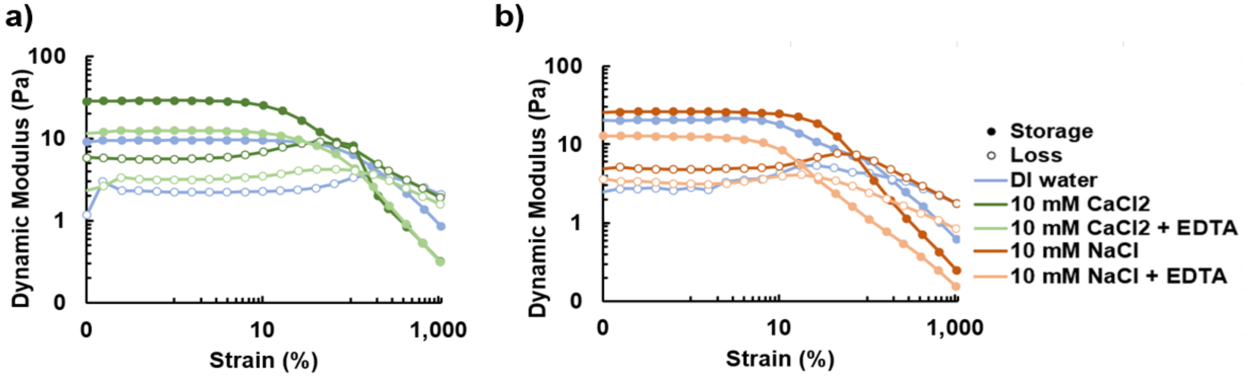

Figure S3: Strain sweep results comparing the addition of a) bivalent ions and b) monvalent ions.

#### 1.4 Dialysis-Based Purification Process

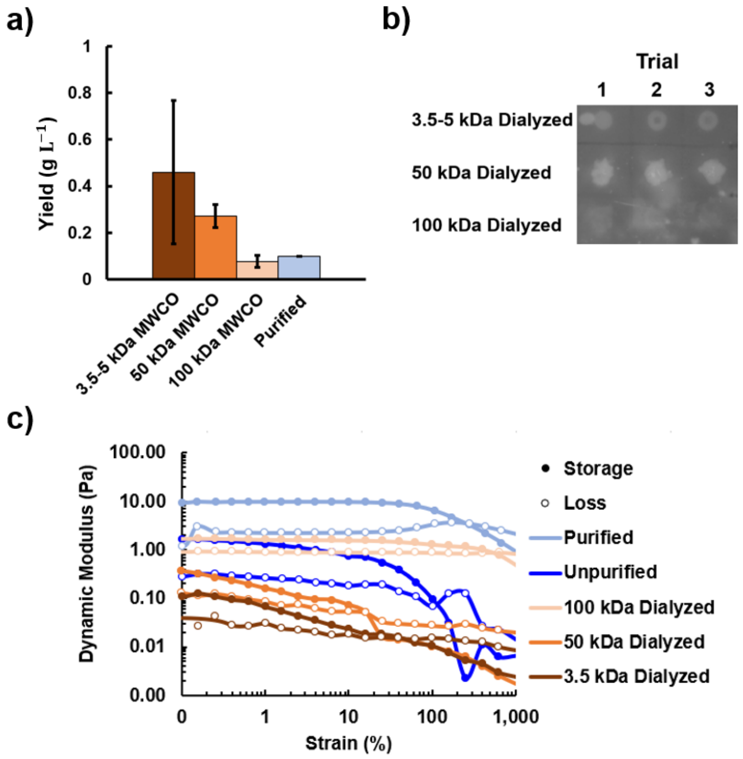

Figure S4: a) Magnified view of sample yields from varying dialysis MWCO values compared to the yield of the full enzymatic purification process. b) A dot blot showing lectin-glycan binding for dialysis groups confirming the presence of polysaccharide c) Strain sweep of samples at various levels of purification.

#### 1.5 The Effect of Ultracentrifugation on Lectin Binding and Rheological Properties

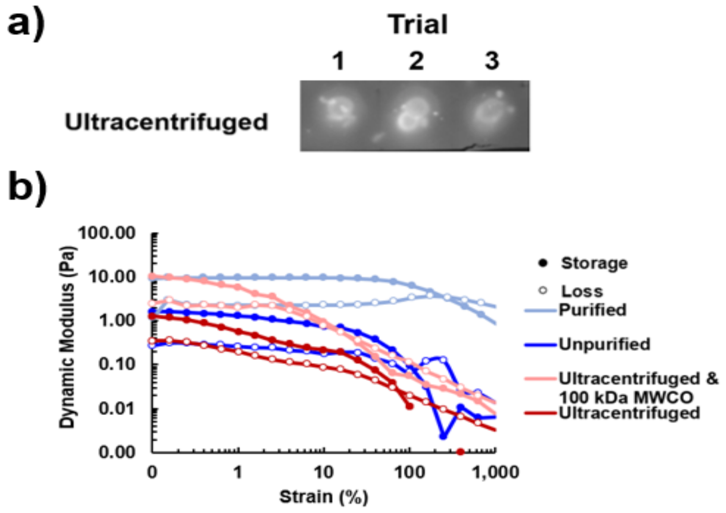

Figure S5: a) A dot blot showing lectin-glycan binding in the sample confirming the presence of polysaccharide post ultracentrifugation. b) Strain sweep of purified, unpurified, and ultracentrifuged samples.

#### 1.6 Rheological Properties of Protein Knockout Material

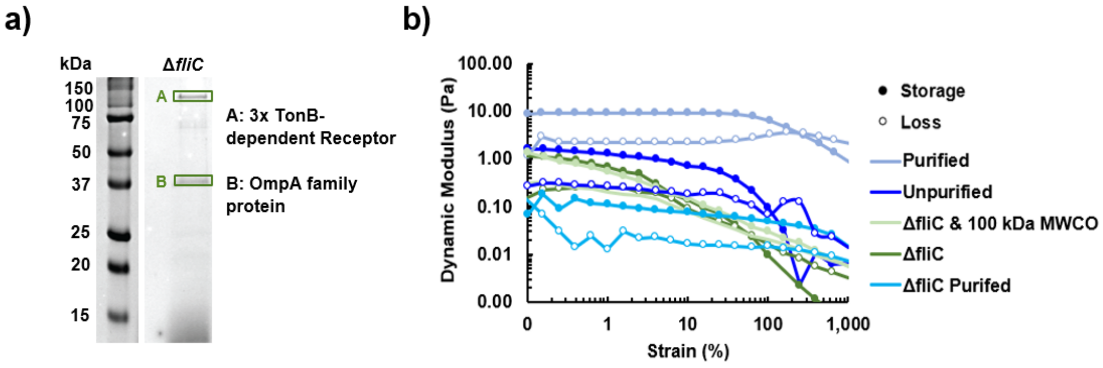

Figure S6: a) Secreted proteins from the Δ*prmJ* Δ*fliC* strain b) Strain sweep of purified, unpurified, and protein knockout material samples.

#### 1.7 Extracellular DNA Analysis

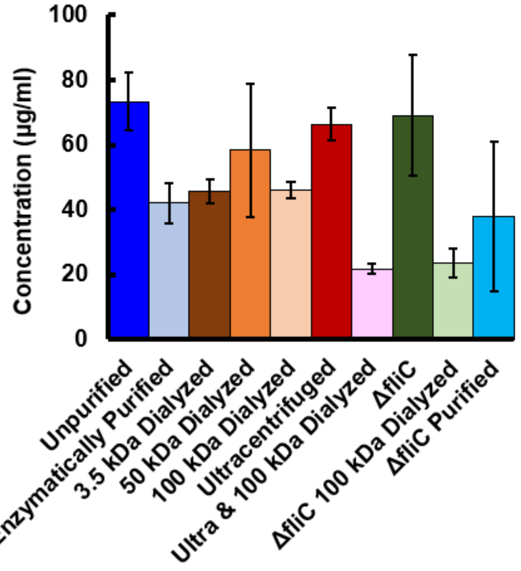

Figure S7: Extracellular DNA abundance in samples tested.

#### 1.8 Thermogravimetric Analysis

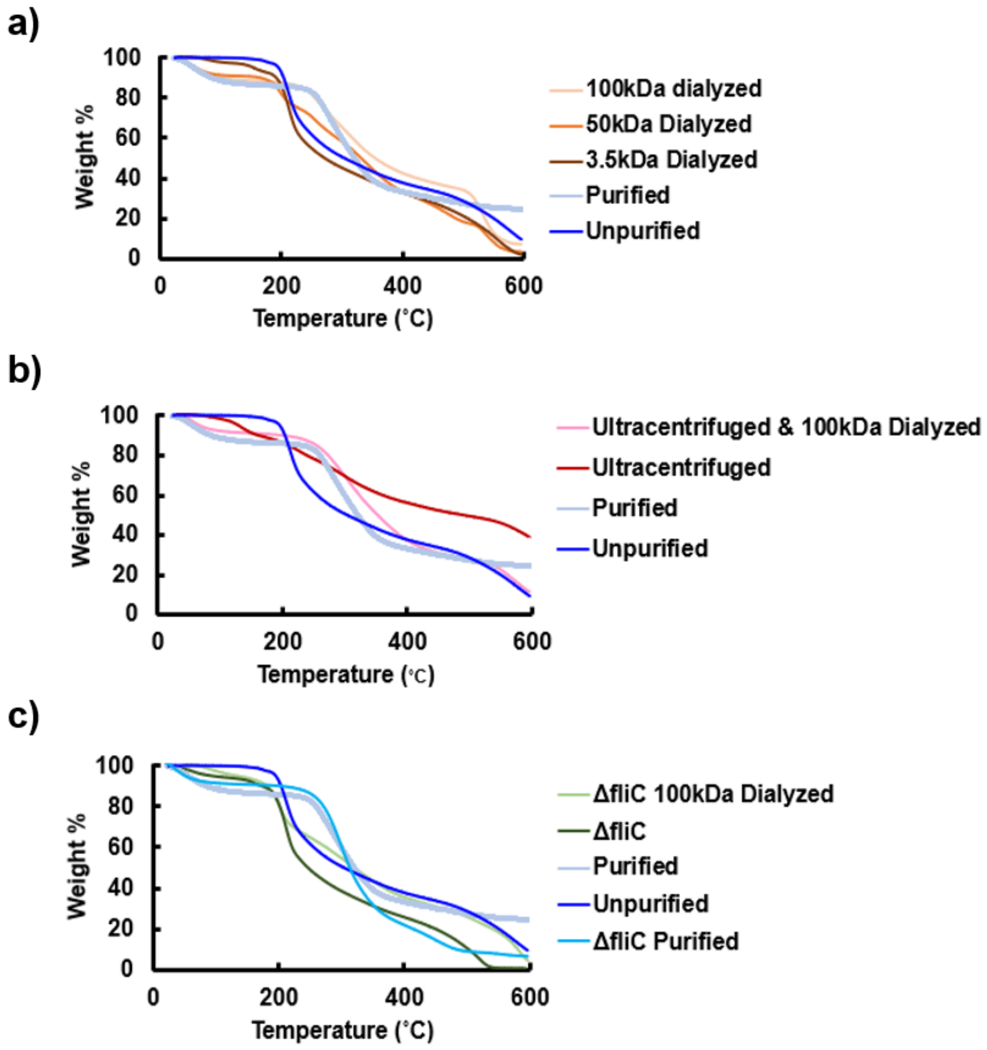

Figure S8: Thermogravimetric analysis for a) dialyzed material, b) ultracentrifuged material and c) protein knockout material.

### 2 Supporting Tables

#### 2.1 Strains Used in this Study

Table S1: Strains used in this Study

| Strain | Organism | Genotype | Description | Source |
|---|---|---|---|---|
| DH173 | *Sphingomonas sp.* LM7 | LM7 | Wild Type | Hershey Lab[1] |
| DH313 | *Sphingomonas sp.* LM7 | $\Delta prmJ$ | In frame deletion of BXU08_RS00395 | Hershey Lab[1] |
| DH1312 | *Sphingomonas sp.* LM7 | $\Delta fliC$ | In frame deletion of BXU08_RS13585 | This work |
| DH1313 | *Sphingomonas sp.* LM7 | $\Delta prmJ \Delta fliC$ | In frame deletion of BXU08_RS13585 in DH313 background | This work |

#### 2.2 Plasmids Used in this Study

Table S2: Plasmids used in this Study

| Plasmid | Description | Antibiotic | Reference |
|---|---|---|---|
| pNPTS138 | Suicide plasmid for deletion in *Sphingomonas sp.* LM7; carries *sacB* for counter-selection | Km | M. Alley[1] |
| pDH298 | To delete *prmJ*; Gibson cloning of fused upstream and downstream regions of BXU08_RS00395 | Km | Hershey Lab[1] |
| pDH1329 | To delete *fliC*; commercial synthesis of fused BXU08_RS13585 upstream and downstream regions, inserted into SpeI/HindIII site of pDH100 | Km | Hershey Lab[1] |
| pDH1331 | To delete *fliC*; commercial synthesis of fused BXU08_RS13585 upstream and downstream regions, inserted into SpeI/HindIII site of pDH100 | Km | This work |

### References

[1] Alexandra G. Goetsch et al. *A novel exopolysaccharide pathway from a freshwater Sphingomonas isolate*. Nov. 4, 2023. DOI: 10.1101/2023.11.03.565537. URL: https://www.biorxiv.org/content/10.1101/2023.11.03.565537v1 (visited on 11/13/2023).
